# JAK2V617F mutant megakaryocytes contribute to hematopoietic aging in a murine model of myeloproliferative neoplasm

**DOI:** 10.1101/2021.08.24.457582

**Authors:** Sandy Lee, Helen Wong, Melissa Castiglione, Malea Murphy, Kenneth Kaushansky, Huichun Zhan

**Author notes:** Correspondence: Huichun Zhan, Division of Hematology-Oncology, Department of Medicine, Stony Brook University, Stony Brook, NY 11794; Northport VA Medical Center, 79 Middleville Road, Northport, NY 11768, USA. **AUTHOR CONTRIBUTION**, S.L. performed various *in vitro* and *in vivo* experiments of the project; H.W. performed/assisted donor chimerism analysis of competitive and serial transplantation experiments; M.C. performed/assisted HSC *in vivo* proliferation assays; M.M. provided technical consultation for whole-mount immunostaining of tibia samples; K.K. reviewed data and revised the manuscript; H. Zhan conceived the projects, analyzed the data, interpreted the results, and wrote the manuscript.

## Abstract

Megakaryocytes (MKs) is an important component of the hematopoietic niche. Abnormal MK hyperplasia is a hallmark feature of myeloproliferative neoplasms (MPNs). The JAK2V617F mutation is present in hematopoietic cells in a majority of patients with MPNs. Using a murine model of MPN in which the human JAK2V617F gene is expressed specifically in the MK lineage, we show that the JAK2V617F-bearing MKs promote hematopoietic stem cell (HSC) aging, manifesting as myeloid-skewed hematopoiesis with an expansion of CD41^+^ HSCs, a reduced engraftment and self-renewal capacity, and a reduced differentiation capacity. HSCs from 2yr old mice with JAK2V617F-bearing MKs were more proliferative and less quiescent than HSCs from age-matched control mice. Examination of the marrow hematopoietic niche reveals that the JAK2V617F-bearing MKs not only have decreased direct interactions with hematopoietic stem/progenitor cells during aging, but also suppress the vascular niche function during aging. Unbiased RNA expression profiling reveals that HSC aging has a profound effect on MK transcriptomic profiles, while targeted cytokine array shows that the JAK2V617F-bearing MKs can alter the hematopoietic niche through increased levels of pro-inflammatory and anti-angiogenic factors. Therefore, as a hematopoietic niche cell, MKs represent an important connection between the extrinsic and intrinsic mechanisms for HSC aging.

**Significance Statement:** The relative contribution of intrinsic and extrinsic mechanisms to HSC aging remains debated. We find that JAK2V617F mutant MKs can accelerate hematopoietic aging both directly (via decreased MK-HSC interaction) and indirectly (via suppressing vascular niche function). We also show that HSC aging has a profound effect on MK function. Our data suggest that, as a hematopoietic niche cell, MKs represent an important connection between HSC-intrinsic and HSC-extrinsic aging mechanisms.

## Introduction

Blood cell production is maintained throughout life by rare multipotent hematopoietic stem cells (HSCs). Dysfunction within the HSC compartment contributes to many age-related diseases including an increased incidence of hematological malignancies in the elderly. HSC aging is characterized by an expansion of phenotypically defined HSCs with impaired functions, such as reduced engraftment and self-renewal capacity, a perturbed state of quiescence, and a skewed differentiation towards the myeloid lineage^1,2^. While these aging-associated HSC functional changes are well established, the molecular and cellular mechanisms that contribute to HSC aging are less well understood. The hematopoietic microenvironment (niche) interacts with HSCs to orchestrate their survival, proliferation, self-renewal, and differentiation. Studies over the past decade suggest that HSC aging is driven by both cell-intrinsic alterations in the stem cells^3–7^, and cell-extrinsic mediators from the aged microenvironment in which the stem cells reside^8–11^. However, the relative contribution of intrinsic and extrinsic mechanisms to HSC aging remains debated. One key question is whether microenvironmental alterations initiate HSC aging or whether aged HSCs cause niche remodeling.

The myeloproliferative neoplasms (MPNs), including polycythemia vera, essential thrombocythemia, and primary myelofibrosis, are clonal stem cell disorders characterized by overproduction of mature blood cells, and increased risk of transformation to acute leukemia or myelofibrosis. An acquired kinase mutation, JAK2V617F, is present in most patients with MPNs and aberrant JAK-STAT signaling plays a central role in these diseases^12^. JAK2V617F is also one of the most frequently mutated genes associated with clonal hematopoiesis of indeterminate potential (CHIP), which is defined as the presence of a somatic mutation in at least 2-4% of blood cells without other hematologic abnormalities. The incidence of both MPNs and CHIP increases significantly with aging^13–18^. These observations indicate a close association between the JAK2V617F mutation and hematopoietic aging. Various murine models of JAK2V617F-positive MPNs were mostly followed for less than 3-9 months^19–30^, providing little to no information of how the JAK2V617F mutation affects HSC aging in MPNs.

Megakaryocytes (MKs) are rare polyploid marrow cells that give rise to blood platelets. MK hyperplasia is a hallmark feature of MPNs^31^ and many MPN-associated genetic mutations/deregulations are preferentially enriched in MKs^32–34^. Very recent evidence has implicated MKs in regulating HSC quiescence and proliferation during both steady-state and stress hematopoiesis, mediated by the many cytokines and extracellular matrix components produced by these cells^35–41^. In contrast to the non-hematopoietic niche cells (e.g. endothelial cells, perivascular stromal cells), niche MKs provide direct feedback to their HSC precursors, many of which are located directly adjacent to MKs *in vivo*.^37,38^ In the present study, we investigated the effects of JAK2V617F-bearing MK niche on HSC aging in a murine model of MPN during a 2-yr follow up.

## Methods

### Experimental mice

JAK2V617F Flip-Flop (FF1) mice (which carry a Cre-inducible human JAK2V617F gene driven by the human JAK2 promoter)^24^ and Pf4-Cre mice (which express Cre under the promoter of platelet factor 4, a MK-specific gene)^42^ were provided by Radek Skoda (University Hospital Basal, Switzerland). The FF1 mice and Pf4-Cre mice were crossed to generate MK lineage-specific human JAK2V617F transgenic mouse line (Pf4-cre^+^FF1^+^, or Pf4^+^FF1^+^)^43–45^. CD45.1+ congenic mice (SJL) were purchased from Taconic Inc. (Albany, NY, USA). Animal experiments were performed in accordance with the Institutional Animal Care and Use Committee guideline.

### Stem cell transplantation assays

For competitive transplantation, 5×10^5^ marrow cells from young (6mo old) or old (2yr old) Pf4-cre control or Pf4^+^FF1^+^ mice (CD45.2) were injected intravenously together with 5×10^5^ marrow cells from 8wk old wild-type mice (CD45.1) into lethally irradiated (950cGy) 8-12wk old wild-type recipient mice (CD45.1). Two independent experiments were performed.

For secondary and tertiary transplantation, primary and secondary recipients were sacrificed at 24wk after transplant and their marrow cells were isolated and pooled. ~5 ×10^6^ marrow cells were transplanted without competitor cells into lethally irradiated wild-type recipients (CD45.1) by intravenous tail vein injection.

### BrdU incorporation analysis

Mice were injected intraperitoneally with a single dose of 5-bromo-2′-deoxyuridine (BrdU; 100 mg/kg body weight) and maintained on 1mg BrdU/ml drinking water for two days. Mice were then euthanized and marrow cells isolated as described above. For analysis of HSC (Lin^-^cKit^+^Sca1^+^CD150^+^CD48^-^) proliferation, Lineage^neg^ (Lin^-^) cells were first enriched using the Lineage Cell Depletion Kit (Miltenyi Biotec) before staining with fluorescent antibodies specific for cell surface HSC markers, followed by fixation and permeabilization using the Cytofix/Cytoperm kit (BD Biosciences, San Jose, CA), DNase digestion (Sigma, St. Louis, MO), and anti-BrdU antibody (Biolegend, San Diego, CA) staining to analyze BrdU incorporation. For analysis of more abundant cell populations, marrow cells were stained with cell surface antibodies, then fixed and stained with anti-BrdU antibody for BrdU incorporation analysis as described above.

### Cell cycle analysis

For HSPC (CD150^+^CD48^-^) cell cycle analysis, marrow cells were first stained with fluorescent antibodies for cell surface HSPC markers, washed, and then stained with Hoechst33342 (10ug/ml) (Sigma) at 37°C in dark for 45 min, followed by staining with Pyronin Y (0.2ug/ml) (Sigma) at 37°C in dark for another 15 min. Cells were kept on ice until flow cytometry analysis on a LSR II (BD biosciences)^45,46^.

### Half bone whole-mount tissue preparation for imaging

Freshly harvested tibias were fixed in 4% paraformaldehyde (PFA) in PBS (Affymetrix) for 6hr at 4°C while rotating. The bones were washed in PBS overnight to remove PFA and cryoprotected in 20% sucrose PBS solution at 4°C. The bones were embedded in OCT (Tissue-Tek) and flash frozen at −80°C. A Leica CM1510S cryostat was used to longitudinally shave the bones until the marrow cavity was exposed. The half tibias were washed in PBS to remove OCT then processed for blocking, staining, clearing, and imaging as below.

### Whole-mount immunostaining

Half tibias were blocked in a PBS buffer containing 10% dimethyl sulfoxide, 0.5% IgePal630 (Sigma Aldrich), 5% donkey serum (Sigma Aldrich), and 0.3 M glycine overnight at room temperature. This blocking buffer was used for all subsequent antibody staining. After blocking, tissues were stained for three days at room temperature with unconjugated goat anti-cKit antibody (R&D Systems) at a 1:250 dilution. Then the tissues were washed multiple times in PBS at room temperature for one day and put into a staining solution containing Alexa Fluor 488 anti-mouse CD41 antibody (clone MWReg30, BioLegend) at a 1:100 dilution and Alexa Fluor 555 donkey anti-goat antibody (ThermoFisher) at a 1:250 dilution for three days, followed by a one-day wash to remove any unbound antibodies. Details of tissue clearing are provided in Supplementary Information^47^.

### Confocal imaging of thick tissue and image analysis

Images of half tibia samples were acquired with Olympus IX81 microscope using 20x objective magnification and Olympus Fluoview FV1000 confocal laser scanning system at 512×512 pixel resolution with 5μm Z-steps. Images were analyzed using Olympus Fluoview Ver.4.2b. We identified HSPCs as having a round morphology and c-kit expression surrounding the cell surface. MKs were distinguished by their size, morphology, and CD41 expression. The number of cKit+ HSPCs adjacent or non-adjacent to CD41+ MKs were imaged and counted in 3 randomly selected bone marrow areas from each sample (n=3 samples in each group). Cells in direct contact or within one HSPC cell distance were considered adjacent.

### VE-cadherin and image analysis

25ug Alexa Fluor 647-conjugated monoclonal antibodies that target mouse VE-cadherin (clone BV13, Biolegend) were injected retro-orbitally into 2yr old Pf4^+^FF1^+^ or control mice under anesthesia^48^. Ten minutes after antibody injection, the mice were euthanized. Tibias were dissected out and washed in PBS. After fixation in 4% PFA for 6hr at 4°C while rotating, the bones were cryoprotected in 20% sucrose, embedded in OCT compound, and snap-frozen. The tibias were cleared using modified Murray’s clear and imaged using Olympus Fluoview FV1000 confocal laser scanning system as described above. Images were analyzed using ImageJ software (National Institute of Health) and VE-cadherin^+^ vascular area was quantified from equal sized 40X stacked images. The sum of analyzed particles was taken from adjustment of the color threshold^49^.

### Statistical analysis

Statistical analysis was performed using Student’s t tests (2 tailed) using Excel software (Microsoft). A p value of less than 0.05 was considered significant. Data are presented as mean ± standard error of the mean (SEM).

Additional methods can be found in Supplementary Methods.

## Results

### The essential thrombocythemia phenotype in the Pf4^+^FF1^+^ mice during a 2-yr follow up

In our previous work, we crossed mice that bear a Cre-inducible human JAK2V617F gene (FF1) with mice that express Cre specifically in the MK lineage (Pf4-Cre) to express JAK2V617F restricted to MK lineage^43–45^. Using this model, we showed that JAK2V617F-bearing MKs caused an essential thrombocythemia phenotype with modest thrombocytosis, splenomegaly, increased marrow MKs, and an expansion of hematopoietic stem/progenitor cells (HSPCs) after 16 wks of age^44^.

The Pf4^+^FF1^+^ mice continued to display modest thrombocytosis compared to age-matched Pf4-cre control mice during long-term follow up. After 1.5 yrs of age, the mice also developed a significant neutrophilia. There was no difference in hemoglobin or total lymphocyte count between Pf4^+^FF1^+^ mice and age-matched control mice (Figure 1A-B). Moderate to severe splenomegaly was present in Pf4^+^FF1^+^ mice compared to control mice and was more prominent during aging (Figure 1C). Quantitative evaluation of the marrow hematopoietic compartment by flow cytometry analysis revealed a significant expansion of CD41^+^ MKs, Lin^-^cKit^+^Sca1^+^ (LSK) HSPCs, and Lin^-^cKit^+^Sca1^+^CD150^+^CD48^-^ HSCs^50^ in both young (~6mo) and old (~2yr) Pf4^+^FF1^+^ mice compared to control mice (Figure 1D-F). There was no difference in total marrow cell counts between Pf4^+^FF1^+^ mice and age-matched control mice (Figure 1G). Histologic examination of marrow and splenic reticulin stained sections did not reveal any significant fibrosis in 2yr old Pf4^+^FF1^+^ mice compared to control mice (Figure 1H-I). Therefore, the Pf4^+^FF1^+^ mice maintained an essential thrombocythemia phenotype during aging with no evidence of transformation to leukemia or myelofibrosis.

**Figure 1.**
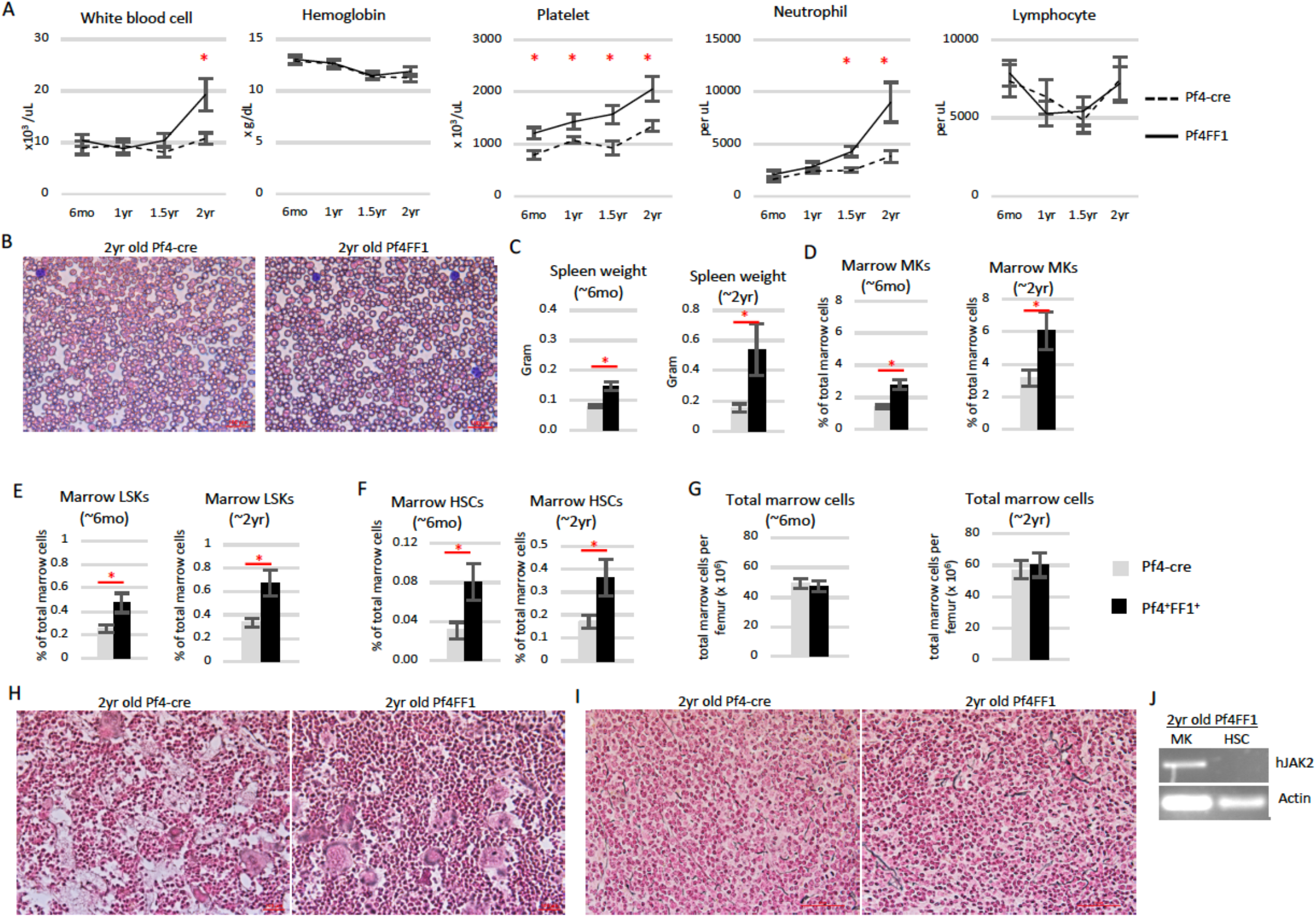
Pf4^+^FF1^+^ mice maintain an essential thrombocythemia phenotype during a 2-yr follow up. (**A**) Peripheral blood cell counts of Pf4^+^FF1^+^ (black line) and Pf4-cre control mice (dotted line). (n=10-12 mice in each group at 6mo, 1yr, and 1.5yr; n=20 mice in each group at 2yr). (**B**) Representative peripheral blood smear of 2yr old Pf4-cre and Pf4^+^FF1^+^ mice (40X magnification) (**C**) Spleen weight in 6mo old (n=10 mice in each group) and 2yr old (n=20 mice in each group) Pf4-cre control and Pf4^+^FF1^+^ mice. (**D-F**) Marrow CD41^+^ MK (D), Lin^-^cKit^+^Sca1^+^ (LSK) HSPCs (E), and Lin^-^cKit^+^Sca1^+^CD150^+^CD48^-^ HSCs (F) frequency of Pf4-cre control and Pf4^+^FF1^+^ mice at 6mo old (n=5 mice in each group) and 2yr old (n=5 mice in each group for D; n=9-10 mice in each group for E-F). (**G**) Total marrow cell numbers per femur in Pf4-cre and Pf4^+^FF1^+^ mice at 6mo old and 2yr old (n=7 mice in each group). (**H-I**) Representative reticulin stain of 2yr old Pf4-cre and Pf4^+^FF1^+^ mice marrow (H) and spleen (I). (**J**) As determined by RT-PCR, human JAK2 was expressed in MK cells, but not in sorted HSCs from 2yr old Pf4^+^FF1^+^ mice. * *P* < 0.05

Previously, we and others used rigorous, sensitive assays to eliminate the possibility that Pf4-cre is expressed in HSCs^37,42,44,51,52^. To be certain that JAK2V617F did not directly influence HSC aging because the Pf4 promoter was ‘leaky’^53–55^, we tested purified HSCs by human JAK2 RT-PCR. Consistent with our previous reports in young (5-6mo) Pf4^+^FF1^+^ mice^44^, human JAK2 gene expression was detected in MKs but not in HSCs in 2yr old Pf4^+^FF1^+^mice (Figure 1J). Similar results were obtained in CD45^+^CD201^+^CD48^-^CD150^+^ (E-SLAM) cells^50,56^, which is a highly purified long-term repopulating HSC population (data not shown).

### Accelerated HSC aging in the Pf4^+^FF1^+^ mice

Consistent with the myeloid-dominant hematopoiesis in peripheral blood of aged Pf4^+^FF1^+^ mice (Figure 1A), myeloid-biased CD41^+^ HSCs^57^ were significantly expanded in 1yr old and 2yr old Pf4^+^FF1^+^ mice compared to age-matched control mice (Figure 2A-B). No difference in CD41^+^ HSC numbers was detected between young (6mo) Pf4^+^FF1^+^ mice and control mice. When HSCs were sorted into individual wells of a 96 well plate containing complete methylcellulose medium, HSCs from old Pf4^+^FF1^+^ mice formed more myeloid colonies than HSCs from old control mice (Figure 2C). These observations indicate a phenotype of chronologically aged HSCs with increased myeloid-biased differentiation.

**Figure 2.**
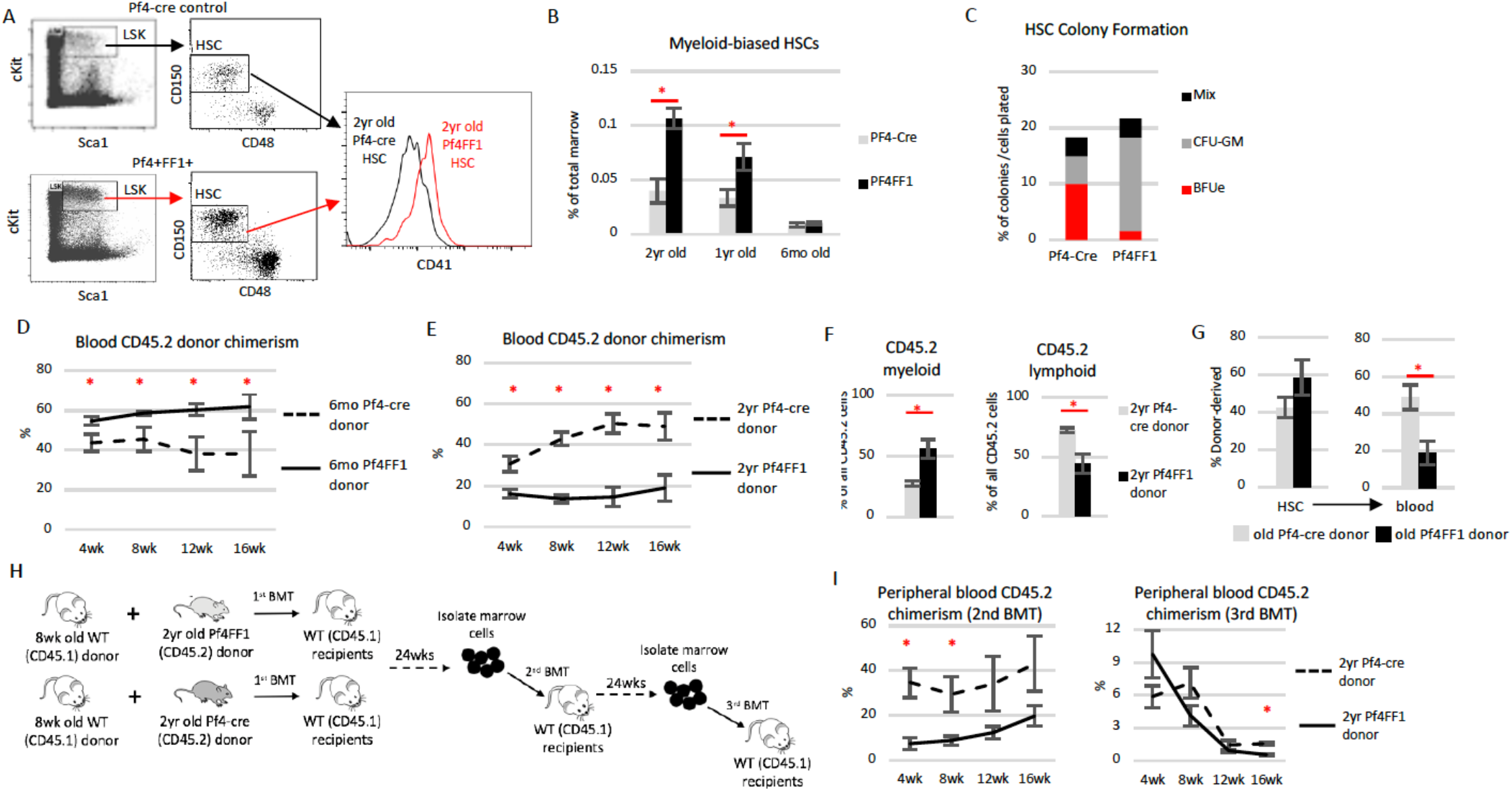
Hallmarks of accelerated HSC aging in the Pf4^+^FF1^+^ mice compared to control mice. (**A-B**) Representative flow cytometry plots showing gating strategy (A) used to measure marrow Lin^-^cKit^+^Sca1^+^CD150^+^CD48^-^CD41^+^ myeloid-biased HSC frequency (B) in 6mo, 1yr, and 2yr old Pf4-cre control and Pf4^+^FF1^+^ mice (6mo and 1yr old: n=3 in each group; 2yr old: n=6 in each group). (**C**) Methylcellulose colony formation: single cells from old Pf4-cre or old Pf4^+^FF1^+^ HSC population were sorted into individual wells of a 96 well plate containing complete methylcellulose medium. After 12-14 days, colonies were scored. The percentage of each type of colony out of the total cells plated is indicated. Data shown is representative from one of two independent experiments of 60 cells each, for a total of 120 cells each group. (**D**) Peripheral blood CD45.2 donor chimerism in recipients of competitive transplantation experiments during which donor marrow cells from 6mo old Pf4-cre or Pf4^+^FF1^+^ mice (CD45.2) were transplanted together with 8wk old CD45.1 wild-type competitor marrow cells into lethally irradiated CD45.1 recipients. (n=8 mice in each group from two independent experiments) (**E**) Peripheral blood CD45.2 donor chimerism in recipients of competitive transplantation experiments during which donor marrow cells from 2yr old Pf4-cre or Pf4^+^FF1^+^ (CD45.2) mice were transplanted together with 8wk old CD45.1 wild-type competitor marrow cells into lethally irradiated CD45.1 recipients. (n=8-9 mice in each group from two independent experiments) (**F**) Frequency of donor-derived myeloid and lymphoid cells in the blood of recipients of 2yr old Pf4-cre or Pf4^+^FF1^+^ marrow donors at 16wks post transplantation. Myeloid cells were defined as CD11b- and/or Gr1-positive events. Lymphoid cells were defined as CD3- or B220-positive events that were negative for CD11b or Gr1. The proportion of myeloid and lymphoid cells from Pf4-cre or Pf4^+^FF1^+^ donor (CD45.2) or wild-type competitor donor (CD45.1) was determined using CD45.2 versus CD45.1 for each cell type. (n=8-9 mice in each group from two independent experiments) (**G**) HSCs from old Pf4^+^FF1^+^ mice are impaired in differentiation compared to HSCs from old Pf4-cre control mice, as shown by their donor-derived marrow HSC chimerism (left) and peripheral blood chimerism (right) (n=5-6 mice in each group). (**H**) Scheme of serial marrow transplantation experiments. (**I**) Peripheral blood CD45.2 donor chimerism in secondary and tertiary transplant recipients after transplantation (n=5 mice in each group).

We then performed two groups of competitive repopulation assays to investigate how the JAK2V617F mutant MK niche might affect HSC engraftment capacity during aging. In the first group, CD45.2 donor marrow cells from 6mo old Pf4^+^FF1^+^ mice or age-matched control mice were injected intravenously together with 8wk old CD45.1 wild-type competitor marrow cells into lethally irradiated (950cGy) CD45.1 recipient mice. During a 16-wk follow up, recipients of Pf4^+^FF1^+^ marrow cells displayed higher peripheral blood donor (CD45.2) chimerism than recipients of the control mouse marrow cells (Figure 2D), consistent with our previous observations^45^. In the second group, CD45.2 donor marrow cells from 2yr old Pf4^+^FF1^+^ mice or age-matched control mice were injected intravenously together with 8wk old CD45.1 wild-type competitor marrow cells into lethally irradiated CD45.1 recipient mice. Recipients of old Pf4^+^FF1^+^ marrow cells displayed a significantly lower peripheral blood CD45.2 chimerism than recipients of the control mice (Figure 2E). In addition, old Pf4^+^FF1^+^ donor marrow cells gave rise to greater numbers of myeloid than lymphoid blood cell output in recipient mice compared to old control donor marrow cells (Figure 2F). These observations suggest that HSCs from old Pf4^+^FF1^+^ mice have reduced engraftment capacity and a skewed differentiation towards the myeloid lineage. We also found that, although peripheral blood CD45.2 donor chimerism was significantly decreased in recipient mice of old Pf4^+^FF1^+^ donor compared to recipients of old control donor, marrow CD45.2 donor-derived HSC chimerism was similar between the two groups (Figure 2G), suggesting a decreased differentiation capacity of the HSCs from old Pf4^+^FF1^+^ mice.

To assess the effects of JAK2V617F mutant MK niche on HSC self-renewal activity, we performed serial transplantation assays using marrow cells from the primary recipients of 2yr old Pf4^+^FF1^+^ mice or Pf4-cre control mice (Figure 2H). Recipients of old Pf4^+^FF1^+^ marrow donor displayed lower peripheral blood CD45.2 donor chimerism than recipients of old Pf4-cre control donor cells (Figure 2I). Taken together, Pf4^+^FF1^+^ mice demonstrated an acceleration of several hallmarks of HSC aging, including an increase in the absolute numbers of HSCs with myeloid-skewed hematopoiesis (and maintenance of this myeloid skewing during marrow transplantation), a reduced engraftment and self-renewal capacity, and a reduced differentiation capacity. Results from the serial transplantation experiments also indicate that the functional decline of HSCs in aged Pf4^+^FF1^+^ mice are HSC-intrinsic and is not reversible.

### JAK2V617F-bearing MK niche promoted HSC proliferation in old Pf4^+^FF1^+^ mice

Cell proliferation is a potent driver of HSC aging^5,58,59^. We examined cell cycle status of CD150^+^CD48^-^ cells, a highly enriched stem/progenitor cell population of which ~ 20% display long-term repopulating capacity^50^, using Hoechst33342 and Pyronin Y staining. Consistent with our previous report^45^, CD150^+^CD48^-^ cells from 6mo old Pf4^+^FF1^+^ mice were more quiescent with a 1.6-fold increase of cells in the G0 phase compared to age-matched control mice (53% vs 34%, *P* = 0.017); in contrast, CD150^+^CD48^-^ cells from 2yr old Pf4^+^FF1^+^ mice were less quiescent with less cells in the G0 phase than wild-type HSPCs from 2yr old Pf4-cre control mice (28% vs 40%, *P* = 0.041) (Figure 3A). We also measured cell proliferation *in vivo* by BrdU labeling. We found that Lin^-^cKit^+^Sca1^+^CD150^+^CD48^-^ HSCs from 2yr old Pf4^+^FF1^+^ mice proliferated more rapidly than cells from 2yr old control mice, while there was no significant difference in HSC proliferation between the young Pf4^+^FF1^+^ mice and control mice (Figure 3B-C). These data suggest that the JAK2V617F mutant MK niche can promote hematopoietic aging in old Pf4^+^FF1^+^ mice by increasing HSC proliferation/cycling.

**Figure 3.**
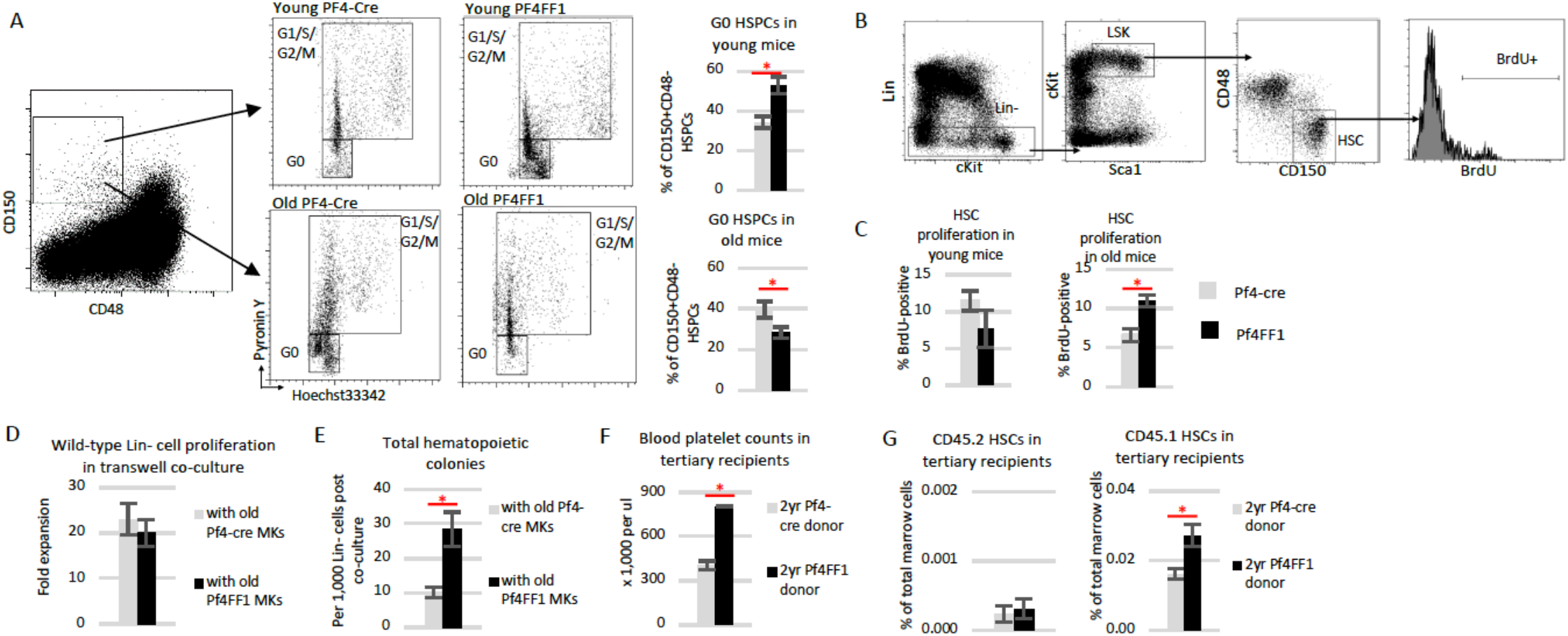
JAK2V617F-bearing MK niche promotes HSC proliferation. (**A**) Representative flow cytometry plots showing gating strategy (left) used to measure G0 cell cycle status of marrow CD150^+^CD48^-^ HSPCs (right) from young (6mo, top) and old (2yr, bottom) Pf4-cre and Pf4^+^FF1^+^ mice measured by Hoechst33342 and Pyronin Y (n=6 mice in each group). (**B-C**) Representative flow cytometry plots showing gating strategy (B) used to measure cell proliferation of Lin^-^cKit^+^Sca1^+^CD150^+^CD48^-^ HSCs from young and old Pf4-cre and Pf4^+^FF1^+^ mice measured by *in vivo* BrdU labeling. (n=4 in young mice group; n=8-9 mice in old mice groups). (**D**) Cell proliferation of wild-type Lin^-^ cells cultured together with wild-type MKs (from 2yr old Pf4-cre control mice) or JAK2V617F mutant MKs (from 2yr old Pf4^+^FF1^+^ mice) in transwells. Cells were cultured in StemSpan serum-free expansion medium containing recombinant mouse SCF (100ng/mL) and recombinant human TPO (100ng/mL). Cell proliferation was shown as fold of expansion which is the ratio of the final cell count to starting cell count. Data are from three independent experiments (with duplicates or triplicates in each experiment). (**E**) Increased colony formation from wild-type Lin^-^ cells after co-culture with JAK2V617F mutant MKs compared to co-culture with wild-type MKs. Data are from three independent experiments (with duplicates in each experiment). (**F**) Peripheral blood platelet counts 16-20wks post transplantation in tertiary recipients of old Pf4^+^FF1^+^ or control donor (n=5 mice in each group). (**F**) Frequency of CD45.2 (left) and CD45.1 (right) marrow HSCs in tertiary recipients of old Pf4^+^FF1^+^ or control donor (n=5 mice in each group).

To further verify what we have observed was not caused by any direct effect of the JAK2V617F mutation on HSC function because the Pf4 promoter was ‘leaky’, we isolated wild-type and JAK2V617F mutant MKs from 2yr old Pf4-cre mice or Pf4^+^FF1^+^ mice and cultured them together with wild-type Lineage^-^ (Lin^-^) HSPCs using a transwell co-culture system. Although there was no difference in the wild-type Lin^-^ cell proliferation between coculture with Pf4-cre MKs and co-culture with Pf4^+^FF1^+^ MKs in serum-free liquid medium, Lin^-^ cells co-cultured with Pf4^+^FF1^+^ MKs generated significantly more hematopoietic progenitor cells on colony formation assay in methylcellulose medium compared to Lin^-^ cells co-cultured with Pf4-cre MKs (Figure 3D-E), suggesting that the mutant MKs can directly promote HSPC proliferation *in vitro* by some secreted factors. These findings prompted us to hypothesize that, in addition to affecting its own precursor HSC function, JAK2V617F mutant MK niche can affect co-existing wild-type competitor HSC function during co-transplantation. To test this hypothesis, we followed recipient mice during the serial transplantation experiments in which old Pf4^+^FF1^+^ or Pf4-cre marrow cells (CD45.2) were co-transplanted with wild-type competitor marrow cells (CD45.1) (see Figure 2H). At 16wk after the tertiary transplantation, recipient mice of old Pf4^+^FF1^+^ donor displayed moderate thrombocytosis compared to control mice (805 vs 406 × 10^9^/L, *P* = 0.0004) (Figure 3F). Quantitative evaluation of their marrow hematopoietic compartment did not reveal any difference in CD45.2 HSC cell frequencies between the two groups; however, CD45.1 HSCs (derived from the co-transplanted wild-type competitor donor) were significantly expanded in the tertiary recipient mice of old Pf4^+^FF1^+^ donor (Figure 3G). Although we do not know whether mutant MK niche would promote co-existing HSC aging due to the short follow up after the tertiary transplantation (i.e. 16-20 wks), these *in vitro* co-cultures and *in vivo* co-transplant experiments demonstrated that the JAK2V617F mutant MK niche can promote co-existing wild-type HSPC expansion.

### Altered hematopoietic microenvironment in old Pf4^+^FF1^+^ mice

HSCs are frequently located adjacent to MKs *in vivo* and MKs can inhibit HSC proliferation and maintain their quiescence^36–38,40^. Decreased interactions between MKs and HSCs have been reported in murine models of aging^11,60^. We examined the spatial relationships between MKs and cKit^+^ HSPCs *in vivo* using whole-mount immunofluorescence staining of thick tibia sections of old Pf4-cre control and Pf4^+^FF1^+^ mice (Figure 4A). We found that cKit^+^ HSPCs were located further from MKs in old Pf4^+^FF1^+^ mice compared to old control mice (Figure 4B-C). These results suggest the possibility that decreased interactions between MKs and HSCs might contribute to the increased HSC proliferation and hematopoietic aging in the Pf4^+^FF1^+^ mice.

**Figure 4.**
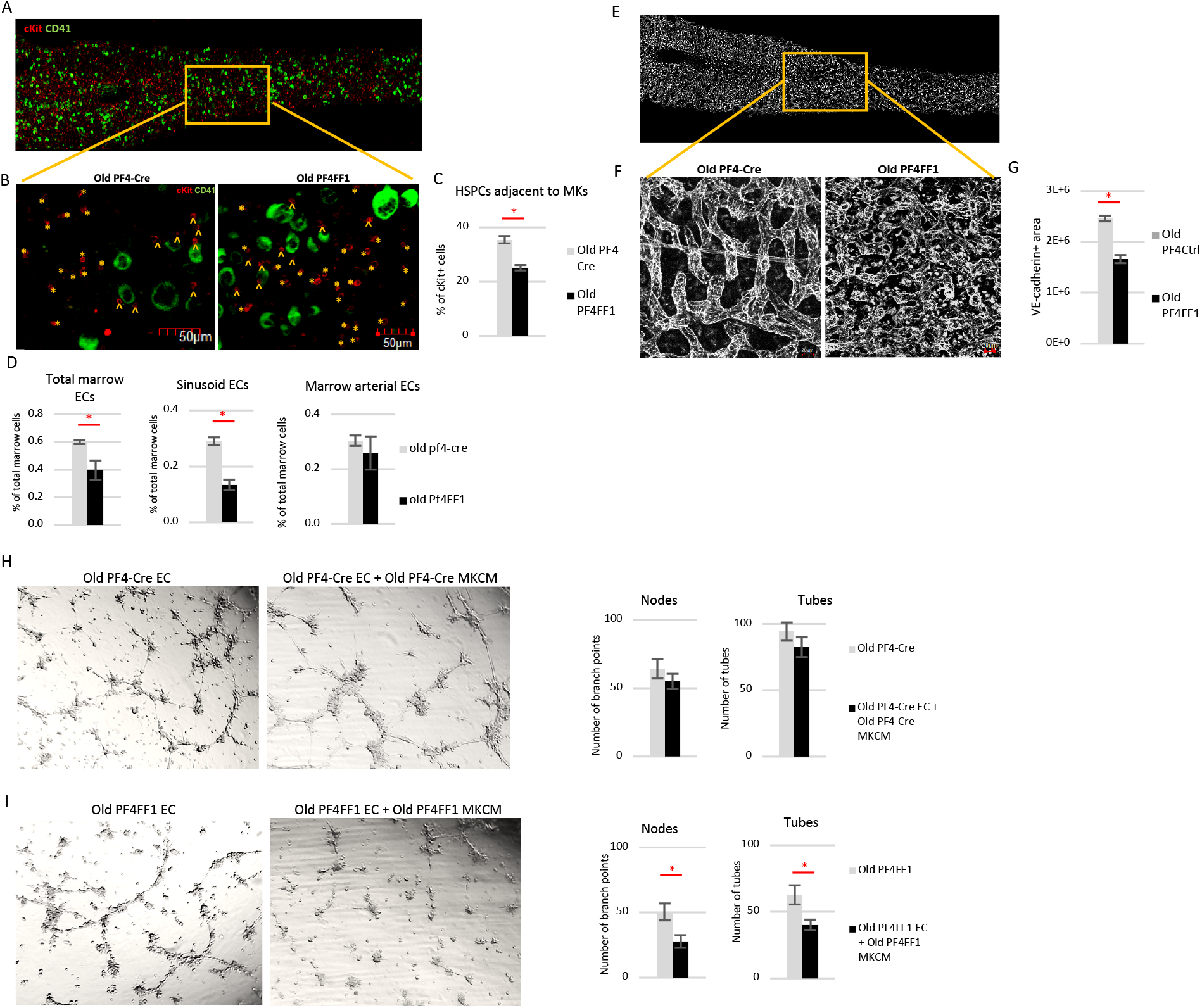
Altered hematopoietic microenvironment in old Pf4^+^FF1^+^ mice. (**A**) Representative whole-mount immunofluorescent staining of thick tibia section showing hematopoietic progenitors (cKit^+^, red) and MKs (CD41^+^, green). Magnification: 20×. (**B-C**) Representative immunofluorescent images (B, magnification 20×) and quantification (C) of cKit^+^ HSPCs adjacent (arrowheads) or non-adjacent (asterisks) to CD41^+^ MKs in the marrow of old Pf4-cre control and Pf4^+^FF1^+^ mice (n=3 mice in each group; a total of 9 non-overlapping areas with ~500 MKs and ~1200 HSPCs were examined for each group). (**D**) Total marrow EC, sinusoidal marrow EC, and arterial marrow EC number in old Pf4-cre control and Pf4^+^FF1^+^ mice (n=3 in each group). (**E**) Representative whole-mount image of thick tibia section, in which vasculature was stained intravenously with anti-VE-cadherin antibody (white). (**F-G**) Representative images (F, magnification 40X) and quantification (G) of VE-cadherin^+^vasculature (white) area in the marrow of old Pf4-Cre control and Pf4^+^FF1^+^ mice (n=2 in each group). For quantification, a total of 12 non-overlapping 500×300 pixel areas at 20× magnification were analyzed for each group. (**H**) (Left) Representative tube formation images of old Pf4-cre lung ECs treated with or without conditioned media of old Pf4-cre MKs. Magnification: 4×. (Right) Quantification of tube formation. Images of tube formation were taken at 4× magnification and quantification was done by counting the number of nodes (or branch points) and tubes in 4 non-overlapping fields. Results are expressed as the mean ± SEM (n=4). Data are from one of two independent experiments that gave similar results. (**I**) (Left) Representative tube formation images of old Pf4^+^FF1^+^ lung ECs treated with or without conditioned media of old Pf4^+^FF1^+^ MKs. Magnification: 4×. (Right) Quantification of tube formation. Data are from one of two independent experiments that gave similar results.

MKs are often located adjacent to marrow sinusoids, a “geography” required for the cells to issue platelets directly into the sinusoidal vascular lumen^61,62^. During aging, the marrow vascular niche exhibits significant morphological and functional changes^9,11,60,63^. Quantitative evaluation by flow cytometry analysis revealed significantly decreased total marrow ECs (CD45^-^CD31^+^) and sinusoidal marrow ECs (CD45^-^CD31^+^Sca1^-^) in aged Pf4^+^FF1^+^ mice compared to control mice, while there was no significant difference in arterial marrow ECs (CD45^-^CD31^+^Sca1^+^) between the two groups (Figure 4D). We examined marrow microvasculature by *in vivo* VE-cadherin labeling and confocal whole-mount imaging of longitudinally shaved tibia of old Pf4^+^FF1^+^ and control mice (Figure 4E-F). Consistent with the flow cytometry analysis findings, vascular area was significantly decreased in the marrow of old Pf4^+^FF1^+^ mice compared to age-matched control mice (Figure 4G).

While ECs have important roles in the regulation of MK maturation and release of platelets^62,64–66^, little is known about the roles of MKs in the regulation of marrow vascular niche, despite MKs representing an important reservoir of bioactive hematopoietic and angiogenic factors. To study the effects of JAK2V617F mutant MKs on EC function *in vitro*, primary murine lung ECs were isolated from old Pf4-cre control mice and Pf4^+^FF1^+^ mice and their tube formation in Matrigel (as a measure of *in vitro* angiogenesis) was assessed. We found that the tube formation of old Pf4-cre lung ECs was not much affected by conditioned medium collected from old Pf4-cre MKs (Figure 4H); in contrast, tube formation of old Pf4^+^FF1^+^ lung ECs were significantly inhibited by conditioned medium collected from old Pf4^+^FF1^+^ MKs (Figure 4I). These findings suggest that the JAK2V617F mutant MK niche not only altered its own interaction with HSCs during aging, but also suppressed the vascular niche function to promote HSC aging.

### Altered inflammatory and angiogenic factors in the JAK2V617F mutant MKs during aging

To further understand the mechanisms by which JAK2V617F mutant MKs promote HSC aging, we performed transcriptomic profiles of wild-type and JAK2V617F mutant MKs from both young (6mo) and old (2yr) Pf4-cre and Pf4^+^FF1^+^ mice. We found that, while young JAK2V617F mutant MKs separated nicely from young wild-type MKs on unsupervised hierarchical clustering analysis, old JAK2V617F mutant MKs were indistinguishable from old wild-type MKs on unsupervised clustering analysis (Figure 5A). Since MKs are constantly generated from HSCs during aging, this finding suggests that HSC aging has a profound effect on MK transcriptomic profiles.

**Figure 5.**
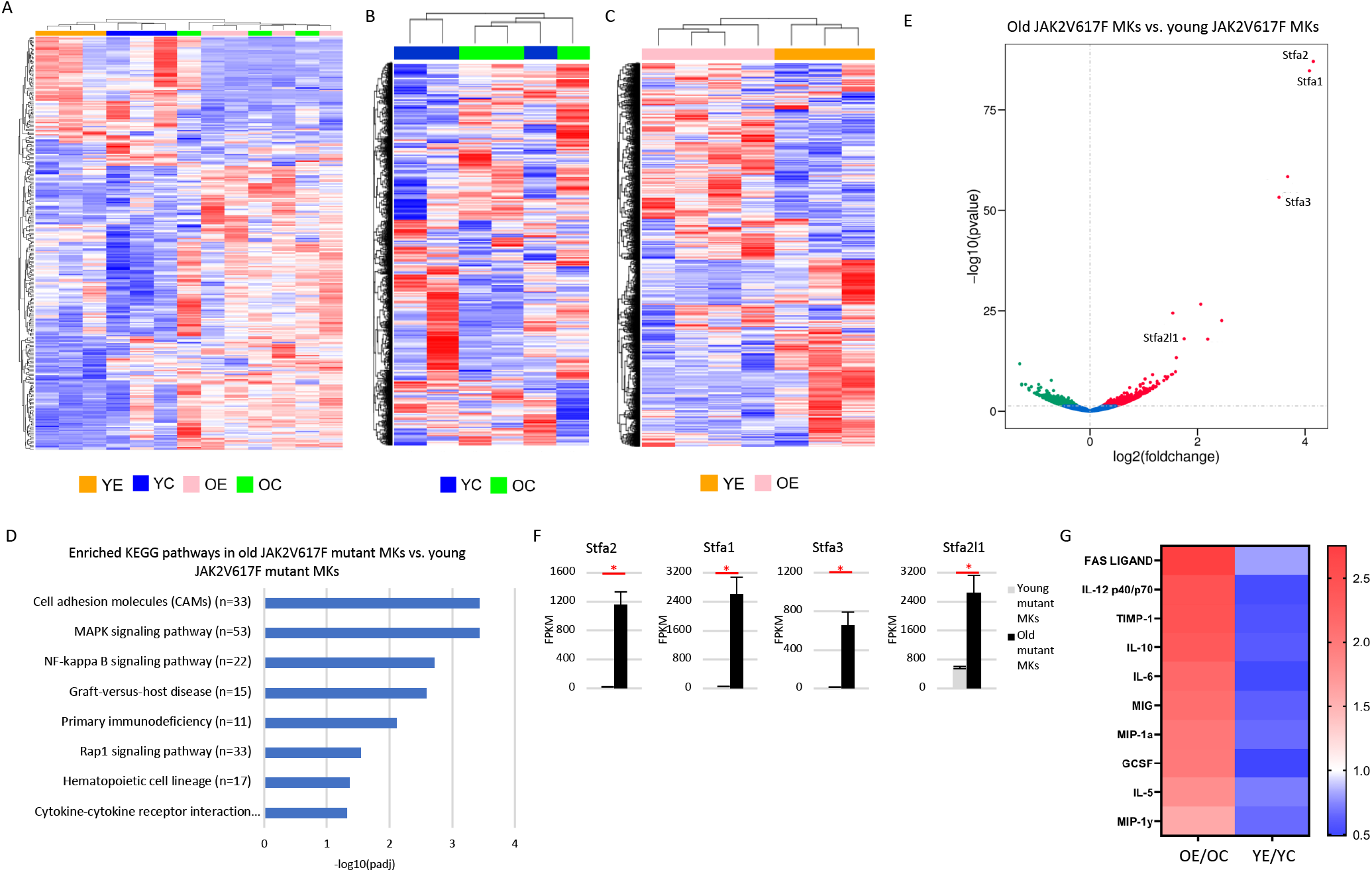
Deregulated MK signaling in aged Pf4^+^FF1^+^ mice. (**A-C**) Unsupervised hierarchical clustering of significantly (p<0.05) deregulated genes in wild-type and JAK2V617F mutant MKs from young and old Pf4-cre control and Pf4^+^FF1^+^ mice (A), in wild-type MKs from young and old Pf4-cre mice (B), and in JAK2V617F mutant MKs from young and old Pf4^+^FF1^+^ mice (C). Young Pf4-cre MKs (YC, n=3), young Pf4^+^FF1^+^ MKs (YE, n=3), old Pf4-cre MKs (OC, n=3), and old Pf4^+^FF1^+^ MKs (OE, n=4). (**D**) Differentially enriched Kyoto lopedia of Genes and Genomes (KEGG) pathways upregulated in old JAK2V617F mutant MKs (from 2’yrold Pf4^+^FF1^+^ mice) compared to young JAK2V617F mutant MKs (from 6mo old Pf4^+^FF1^+^ mice). Adjusted *P* values are plotted as the negative of their logarithm. (**E**) Volcano plot of differentially expressed genes between JAK2V617F mutant MKs from young and old Pf4^+^FF1^+^ mice. (**F**) Stefin A family gene transcript levels in JAK2V617F mutant MKs from young and old Pf4^+^FF1^+^ mice by RNA-seq analysis (n = 3-4). (**G**) Heatmap summary of selected cytokine array data showing factors that were significantly up-regulated in old JAK2V617F mutant MKs (left) but down-regulated in young mutant MKs (right) compared to aged-matched wild-type control MKs. Pooled MK cell lysate from young Pf4-cre control mice (n=4), young Pf4^+^FF1^+^ mice (n=4), old Pf4-cre control mice (n=4), and old Pf4^+^FF1^+^ mice (n=4) were used in the array.

Our previous works and current study have shown that the effect of JAK2V617F mutant MK niche on HSC function is “bimodal” during aging: in young mice, mutant MK niche induces HSC quiescence with increased repopulating capacity and stimulates EC tube formation^44,45^; in old mice, mutant MK niche promotes HSC proliferation with a reduced engraftment and self-renewal capacity and inhibits EC tube formation (Figures 2 and 4). These observations prompted us to examine how the JAK2V617F mutation affects MK transcriptome during aging. While young and old wild-type MK gene expression profiles were mostly indistinguishable on unsupervised clustering analysis (Figure 5B), old JAK2V617F mutant MKs were very different from young JAK2V617F mutant MKs (Figure 5C). Dysregulated pathways in cell adhesion molecules, MAPK signaling, NF-kappa B signaling, hematopoietic cell lineage, and cytokine-cytokine receptor interaction were highly upregulated in old JAK2V617F mutant MKs compared to young JAK2V617F mutant MKs (Figure 5D). Stefin A family of genes (Stfa1/Stfa2/Stfa3/Stfa2l1), which are known cathepsin inhibitors^67–69^, are the most upregulated in old JAK2V617F mutant MKs compared to young mutant MKs, suggesting a disruption of the bone marrow niche in old Pf4^+^FF1^+^ mice (Figure 5E-F).

In order to identify MK proteins that may have contributed to HSC aging in the Pf4^+^FF1^+^ mice, we performed targeted cytokine arrays on wild-type and JAK2V617F mutant MKs from both young (6mo) and old (2yr) Pf4-cre and Pf4^+^FF1^+^ mice. We focused on the difference between young and old JAK2V617F mutant MKs and identified 10 factors (FAS ligand, IL-12, tissue inhibitor of metalloproteinases-1 (TIMP-1), IL-10, IL-6, MIG, macrophage inflammatory protein 1α (MIP-1α), GCSF, IL-5, MIP-1γ) produced in increased amounts in old mutant MKs compared to old wild-type MKs but were decreased in young MKs compared to young wild-type MKs (Figure 5G). Many of these factors are involved in inflammation (e.g. IL-6^70^, IL-12^71^, MIP-1α^72^), angiogenesis (e.g. IL-12^73,74^, IL-10^75,76^), extracellular matrix remodeling (TIMP-1^77^), normal or neoplastic hematopoiesis (e.g. IL-6^11,78,79^, IL-12^80^, G-CSF^81^). These results suggest that the JAK2V617F-bearing MKs can alter the hematopoietic niche (e.g. increased inflammation, decreased angiogenesis) to accelerate HSC aging.

## Discussion

The relative contribution of intrinsic and extrinsic mechanisms to HSC aging remains debated. Results from this study support that, as a hematopoietic niche cell, MKs represent an important connection between the extrinsic and intrinsic mechanisms for HSC aging in MPNs — the JAK2V617F-bearing MKs can alter the hematopoietic niche to accelerate HSC aging, and HSC aging in turn can profoundly remodel the niche e.g. by affecting MK transcriptomics. In addition, we found that the JAK2V617F mutant MK niche not only can promote HSC aging directly via both cell-cell interaction and various secreted factors, but also can inhibit/disrupt the vascular niche to promote HSC aging indirectly. Whether the MK niche function can be harnessed to prevent mutant clone expansion and disease evolution in MPNs will require careful study.

Recently, using mT/mG reporter mice, Mansier et al. reported that the Pf4 promoter could induce recombination in a small subset of HSCs^55^. In our study, we used the same Pf4-cre mice^42^ but a different transgenic JAK2V617F mice^24^. Unlike the JAK2V617F knock-in mouse used by Mansier et al. in which the mice developed a PV-like phenotype at 10wks of age, our Pf4^+^FF1^+^ mice maintained an essential thrombocythemia phenotype during ~2yr follow up. We checked JAK2V617F gene expression by a sensitive PCR assay^44^ and did not detect any leakiness of the Pf4 promoter in either Lin^-^cKit^+^Sca1^+^CD150^+^CD48^-^ HSCs (Figure 1J) or CD45^+^CD201^+^CD48^-^CD150^+^ HSCs. Our *in vitro* co-culture and *in vivo* co-transplantation assays provided further evidence that mutant MKs affected wild-type HSC function directly (Figure 3D-G). The cause(s) for these differing results is not clear, although it is very likely that the aberrant MK development in JAK2V617F-positive MPNs may have contributed to different regulation of the Pf4 promoter in different murine models.

An increase in CD41^+^ HSCs has recently been reported in both JAK2V617F-positive murine models and patients with MPNs^82^, though how these CD41^+^ HSC population expand and whether they promote MPN stem cell exhaustion remain unclear. We found that, while there was no expansion of CD41^+^ HSCs in young (6mo) Pf4^+^FF1^+^ mice, these myeloid-biased HSCs emerged faster in Pf4^+^FF1^+^ mice during aging and were significantly expanded in 1yr- and 2yr-old Pf4^+^FF1^+^ mice (Figure 2B). These findings suggest that the JAK2V617F-bearing MK niche can promote the expansion of these myeloid-biased CD41^+^ HSC as part of an accelerated hematopoietic aging process. We also showed that, while JAK2V617F mutant MKs promoted its precursor HSC aging, they also expanded the surrounding wild-type HSCs during serial competitive transplantations, suggesting that mutant MK niche may play important roles in mutant and wild-type cell competition as well as clonal evolution in MPNs.

## ACKNOWLEDGEMENTS

This research was supported by the National Heart, Lung, and Blood Institute grant NIH R01 HL134970 (H.Z.), VA Career Development Award BX001559 (HZ), and VA Merit Award BX003947 (H.Z.).

## References

1. de Haan G, Lazare SS. Aging of hematopoietic stem cells. Blood 2018;131:479–87.

2. Verovskaya EV, Dellorusso PV, Passegue E. Losing Sense of Self and Surroundings: Hematopoietic Stem Cell Aging and Leukemic Transformation. Trends Mol Med 2019;25:494–515.

3. Rossi DJ, Bryder D, Zahn JM, et al. Cell intrinsic alterations underlie hematopoietic stem cell aging. Proc Natl Acad Sci U S A 2005;102:9194–9.

4. Chambers SM, Shaw CA, Gatza C, Fisk CJ, Donehower LA, Goodell MA. Aging hematopoietic stem cells decline in function and exhibit epigenetic dysregulation. PLoS Biol 2007;5:e201.

5. Flach J, Bakker ST, Mohrin M, et al. Replication stress is a potent driver of functional decline in ageing haematopoietic stem cells. Nature 2014;512:198–202.

6. Ho TT, Warr MR, Adelman ER, et al. Autophagy maintains the metabolism and function of young and old stem cells. Nature 2017;543:205–10.

7. Dykstra B, Olthof S, Schreuder J, Ritsema M, de Haan G. Clonal analysis reveals multiple functional defects of aged murine hematopoietic stem cells. J Exp Med 2011;208:2691–703.

8. Ergen AV, Boles NC, Goodell MA. Rantes/Ccl5 influences hematopoietic stem cell subtypes and causes myeloid skewing. Blood 2012;119:2500–9.

9. Maryanovich M, Zahalka AH, Pierce H, et al. Adrenergic nerve degeneration in bone marrow drives aging of the hematopoietic stem cell niche. Nat Med 2018;24:782–91.

10. Poulos MG, Ramalingam P, Gutkin MC, et al. Endothelial transplantation rejuvenates aged hematopoietic stem cell function. J Clin Invest 2017;127:4163–78.

11. Ho YH, Del Toro R, Rivera-Torres J, et al. Remodeling of Bone Marrow Hematopoietic Stem Cell Niches Promotes Myeloid Cell Expansion during Premature or Physiological Aging. Cell Stem Cell 2019;25:407–18 e6.

12. Nangalia J, Green AR. Myeloproliferative neoplasms: from origins to outcomes. Hematology Am Soc Hematol Educ Program 2017;2017:470–9.

13. Spivak JL. Myeloproliferative Neoplasms. N Engl J Med 2017;376:2168–81.

14. Xie M, Lu C, Wang J, et al. Age-related mutations associated with clonal hematopoietic expansion and malignancies. Nat Med 2014;20:1472–8.

15. Genovese G, Kahler AK, Handsaker RE, et al. Clonal hematopoiesis and blood-cancer risk inferred from blood DNA sequence. N Engl J Med 2014;371:2477–87.

16. Jaiswal S, Fontanillas P, Flannick J, et al. Age-related clonal hematopoiesis associated with adverse outcomes. N Engl J Med 2014;371:2488–98.

17. McKerrell T, Park N, Moreno T, et al. Leukemia-associated somatic mutations drive distinct patterns of age-related clonal hemopoiesis. Cell Rep 2015;10:1239–45.

18. Jaiswal S, Ebert BL. Clonal hematopoiesis in human aging and disease. Science 2019;366.

19. Bumm TG, Elsea C, Corbin AS, et al. Characterization of murine JAK2V617F-positive myeloproliferative disease. Cancer Res 2006;66:11156–65.

20. Lacout C, Pisani DF, Tulliez M, Gachelin FM, Vainchenker W, Villeval JL. JAK2V617F expression in murine hematopoietic cells leads to MPD mimicking human PV with secondary myelofibrosis. Blood 2006;108:1652–60.

21. Wernig G, Mercher T, Okabe R, Levine RL, Lee BH, Gilliland DG. Expression of Jak2V617F causes a polycythemia vera-like disease with associated myelofibrosis in a murine bone marrow transplant model. Blood 2006;107:4274–81.

22. Zaleskas VM, Krause DS, Lazarides K, et al. Molecular pathogenesis and therapy of polycythemia induced in mice by JAK2 V617F. PLoS One 2006;1:e18.

23. Shide K, Shimoda HK, Kumano T, et al. Development of ET, primary myelofibrosis and PV in mice expressing JAK2 V617F. Leukemia 2008;22:87–95.

24. Tiedt R, Hao-Shen H, Sobas MA, et al. Ratio of mutant JAK2-V617F to wild-type Jak2 determines the MPD phenotypes in transgenic mice. Blood 2008;111:3931–40.

25. Xing S, Wanting TH, Zhao W, et al. Transgenic expression of JAK2V617F causes myeloproliferative disorders in mice. Blood 2008;111:5109–17.

26. Akada H, Yan D, Zou H, Fiering S, Hutchison RE, Mohi MG. Conditional expression of heterozygous or homozygous Jak2V617F from its endogenous promoter induces a polycythemia vera-like disease. Blood 2010;115:3589–97.

27. Mullally A, Lane SW, Ball B, et al. Physiological Jak2V617F expression causes a lethal myeloproliferative neoplasm with differential effects on hematopoietic stem and progenitor cells. Cancer Cell 2010;17:584–96.

28. Marty C, Lacout C, Martin A, et al. Myeloproliferative neoplasm induced by constitutive expression of JAK2V617F in knock-in mice. Blood 2010;116:783–7.

29. Li J, Spensberger D, Ahn JS, et al. JAK2 V617F impairs hematopoietic stem cell function in a conditional knock-in mouse model of JAK2 V617F-positive essential thrombocythemia. Blood 2010;116:1528–38.

30. Li J, Kent DG, Godfrey AL, et al. JAK2V617F homozygosity drives a phenotypic switch in myeloproliferative neoplasms, but is insufficient to sustain disease. Blood 2014;123:3139–51.

31. Ciurea SO, Merchant D, Mahmud N, et al. Pivotal contributions of megakaryocytes to the biology of idiopathic myelofibrosis. Blood 2007;110:986–93.

32. Vannucchi AM, Rotunno G, Bartalucci N, et al. Calreticulin mutation-specific immunostaining in myeloproliferative neoplasms: pathogenetic insight and diagnostic value. Leukemia 2014;28:1811–8.

33. Kollmann K, Warsch W, Gonzalez-Arias C, et al. A novel signalling screen demonstrates that CALR mutations activate essential MAPK signalling and facilitate megakaryocyte differentiation. Leukemia 2017;31:934–44.

34. Prestipino A, Emhardt AJ, Aumann K, et al. Oncogenic JAK2(V617F) causes PD-L1 expression, mediating immune escape in myeloproliferative neoplasms. Sci Transl Med 2018;10.

35. Zhao M, Ross JT, Itkin T, et al. FGF signaling facilitates postinjury recovery of mouse hematopoietic system. Blood 2012;120:1831–42.

36. Heazlewood SY, Neaves RJ, Williams B, Haylock DN, Adams TE, Nilsson SK. Megakaryocytes co-localise with hemopoietic stem cells and release cytokines that up-regulate stem cell proliferation. Stem Cell Res 2013;11:782–92.

37. Zhao M, Perry JM, Marshall H, et al. Megakaryocytes maintain homeostatic quiescence and promote post-injury regeneration of hematopoietic stem cells. Nat Med 2014;20:1321–6.

38. Bruns I, Lucas D, Pinho S, et al. Megakaryocytes regulate hematopoietic stem cell quiescence through CXCL4 secretion. Nat Med 2014;20:1315–20.

39. Malara A, Currao M, Gruppi C, et al. Megakaryocytes contribute to the bone marrow-matrix environment by expressing fibronectin, type IV collagen, and laminin. Stem Cells 2014;32:926–37.

40. Nakamura-Ishizu A, Takubo K, Fujioka M, Suda T. Megakaryocytes are essential for HSC quiescence through the production of thrombopoietin. Biochem Biophys Res Commun 2014;454:353–7.

41. Nakamura-Ishizu A, Takubo K, Kobayashi H, Suzuki-Inoue K, Suda T. CLEC-2 in megakaryocytes is critical for maintenance of hematopoietic stem cells in the bone marrow. J Exp Med 2015;212:2133–46.

42. Tiedt R, Schomber T, Hao-Shen H, Skoda RC. Pf4-Cre transgenic mice allow the generation of lineage-restricted gene knockouts for studying megakaryocyte and platelet function in vivo. Blood 2007;109:1503–6.

43. Etheridge SL, Roh ME, Cosgrove ME, et al. JAK2V617F-positive endothelial cells contribute to clotting abnormalities in myeloproliferative neoplasms. Proc Natl Acad Sci U S A 2014;111:2295–300.

44. Zhan H, Ma Y, Lin CH, Kaushansky K. JAK2(V617F)-mutant megakaryocytes contribute to hematopoietic stem/progenitor cell expansion in a model of murine myeloproliferation. Leukemia 2016;30:2332–41.

45. Zhang Y, Lin CHS, Kaushansky K, Zhan H. JAK2V617F Megakaryocytes Promote Hematopoietic Stem/Progenitor Cell Expansion in Mice Through Thrombopoietin/MPL Signaling. Stem Cells 2018;36:1676–84.

46. Shapiro HM. Flow cytometric estimation of DNA and RNA content in intact cells stained with Hoechst 33342 and pyronin Y. Cytometry 1981;2:143–50.

47. Acar M, Kocherlakota KS, Murphy MM, et al. Deep imaging of bone marrow shows non-dividing stem cells are mainly perisinusoidal. Nature 2015;526:126–30.

48. Guo P, Poulos MG, Palikuqi B, et al. Endothelial jagged-2 sustains hematopoietic stem and progenitor reconstitution after myelosuppression. J Clin Invest 2017;127:4242–56.

49. Chen Q, Liu Y, Jeong HW, et al. Apelin(+) Endothelial Niche Cells Control Hematopoiesis and Mediate Vascular Regeneration after Myeloablative Injury. Cell Stem Cell 2019;25:768–83 e6.

50. Kiel MJ, Yilmaz OH, Iwashita T, Yilmaz OH, Terhorst C, Morrison SJ. SLAM family receptors distinguish hematopoietic stem and progenitor cells and reveal endothelial niches for stem cells. Cell 2005;121:1109–21.

51. Ng AP, Kauppi M, Metcalf D, et al. Mpl expression on megakaryocytes and platelets is dispensable for thrombopoiesis but essential to prevent myeloproliferation. Proc Natl Acad Sci U S A 2014;111:5884–9.

52. Woods B, Chen W, Chiu S, et al. Activation of JAK/STAT Signaling in Megakaryocytes Sustains Myeloproliferation In Vivo. Clin Cancer Res 2019;25:5901–12.

53. Chagraoui H, Kassouf M, Banerjee S, et al. SCL-mediated regulation of the cell-cycle regulator p21 is critical for murine megakaryopoiesis. Blood 2011;118:723–35.

54. Calaminus SD, Guitart AV, Sinclair A, et al. Lineage tracing of Pf4-Cre marks hematopoietic stem cells and their progeny. PLoS One 2012;7:e51361.

55. Mansier O, Kilani B, Guitart AV, et al. Description of a knock-in mouse model of JAK2V617F MPN emerging from a minority of mutated hematopoietic stem cells. Blood 2019;134:2383–7.

56. Kent DG, Copley MR, Benz C, et al. Prospective isolation and molecular characterization of hematopoietic stem cells with durable self-renewal potential. Blood 2009;113:6342–50.

57. Gekas C, Graf T. CD41 expression marks myeloid-biased adult hematopoietic stem cells and increases with age. Blood 2013;121:4463–72.

58. Beerman I, Bock C, Garrison BS, et al. Proliferation-dependent alterations of the DNA methylation landscape underlie hematopoietic stem cell aging. Cell Stem Cell 2013;12:413–25.

59. Kirschner K, Chandra T, Kiselev V, et al. Proliferation Drives Aging-Related Functional Decline in a Subpopulation of the Hematopoietic Stem Cell Compartment. Cell Rep 2017;19:1503–11.

60. Sacma M, Pospiech J, Bogeska R, et al. Haematopoietic stem cells in perisinusoidal niches are protected from ageing. Nat Cell Biol 2019;21:1309–20.

61. Junt T, Schulze H, Chen Z, et al. Dynamic visualization of thrombopoiesis within bone marrow. Science 2007;317:1767–70.

62. Avecilla ST, Hattori K, Heissig B, et al. Chemokine-mediated interaction of hematopoietic progenitors with the bone marrow vascular niche is required for thrombopoiesis. Nat Med 2004;10:64–71.

63. Kusumbe AP, Ramasamy SK, Itkin T, et al. Age-dependent modulation of vascular niches for haematopoietic stem cells. Nature 2016;532:380–4.

64. Hamada T, Mohle R, Hesselgesser J, et al. Transendothelial migration of megakaryocytes in response to stromal cell-derived factor 1 (SDF-1) enhances platelet formation. J Exp Med 1998;188:539–48.

65. Rafii S, Shapiro F, Pettengell R, et al. Human bone marrow microvascular endothelial cells support long-term proliferation and differentiation of myeloid and megakaryocytic progenitors. Blood 1995;86:3353–63.

66. Kong Y, Hu Y, Zhang XH, et al. Association between an impaired bone marrow vascular microenvironment and prolonged isolated thrombocytopenia after allogeneic hematopoietic stem cell transplantation. Biol Blood Marrow Transplant 2014;20:1190–7.

67. Staudt ND, Aicher WK, Kalbacher H, et al. Cathepsin X is secreted by human osteoblasts, digests CXCL-12 and impairs adhesion of hematopoietic stem and progenitor cells to osteoblasts. Haematologica 2010;95:1452–60.

68. Grzonka Z, Jankowska E, Kasprzykowski F, et al. Structural studies of cysteine proteases and their inhibitors. Acta Biochim Pol 2001;48:1–20.

69. Mezzapesa A, Bastelica D, Crescence L, et al. Increased levels of the megakaryocyte and platelet expressed cysteine proteases stefin A and cystatin A prevent thrombosis. Sci Rep 2019;9:9631.

70. Tanaka T, Narazaki M, Kishimoto T. IL-6 in inflammation, immunity, and disease. Cold Spring Harb Perspect Biol 2014;6:a016295.

71. Trinchieri G. Interleukin-12: a proinflammatory cytokine with immunoregulatory functions that bridge innate resistance and antigen-specific adaptive immunity. Annu Rev Immunol 1995;13:251–76.

72. Cook DN, Beck MA, Coffman TM, et al. Requirement of MIP-1 alpha for an inflammatory response to viral infection. Science 1995;269:1583–5.

73. Sgadari C, Angiolillo AL, Tosato G. Inhibition of angiogenesis by interleukin-12 is mediated by the interferon-inducible protein 10. Blood 1996;87:3877–82.

74. Strasly M, Cavallo F, Geuna M, et al. IL-12 inhibition of endothelial cell functions and angiogenesis depends on lymphocyte-endothelial cell cross-talk. J Immunol 2001;166:3890–9.

75. Silvestre JS, Mallat Z, Duriez M, et al. Antiangiogenic effect of interleukin-10 in ischemia-induced angiogenesis in mice hindlimb. Circ Res 2000;87:448–52.

76. Kohno T, Mizukami H, Suzuki M, et al. Interleukin-10-mediated inhibition of angiogenesis and tumor growth in mice bearing VEGF-producing ovarian cancer. Cancer Res 2003;63:5091–4.

77. Brew K, Dinakarpandian D, Nagase H. Tissue inhibitors of metalloproteinases: evolution, structure and function. Biochim Biophys Acta 2000;1477:267–83.

78. Reynaud D, Pietras E, Barry-Holson K, et al. IL-6 controls leukemic multipotent progenitor cell fate and contributes to chronic myelogenous leukemia development. Cancer Cell 2011;20:661–73.

79. Zhang TY, Dutta R, Benard B, Zhao F, Yin R, Majeti R. IL-6 blockade reverses bone marrow failure induced by human acute myeloid leukemia. Sci Transl Med 2020;12.

80. Eng VM, Car BD, Schnyder B, et al. The stimulatory effects of interleukin (IL)-12 on hematopoiesis are antagonized by IL-12-induced interferon gamma in vivo. J Exp Med 1995;181:1893–8.

81. Greenbaum AM, Link DC. Mechanisms of G-CSF-mediated hematopoietic stem and progenitor mobilization. Leukemia 2011;25:211–7.

82. Rao TN, Hansen N, Stetka J, et al. JAK2-V617F and interferon-alpha induce megakaryocyte-biased stem cells characterized by decreased long-term functionality. Blood 2021;137:2139–51.

