## Supplemental info for "JAK2V617F mutant megakaryocytes contribute to hematopoietic aging in a murine model of myeloproliferative neoplasm"

### Materials and Methods

*Marrow cell isolation*

Murine femurs and tibias were first harvested and cleaned thoroughly. Marrow cells were flushed into PBS with 2% fetal bovine serum using a 25G needle and syringe. Remaining bones were crushed with a mortar and pestle followed by enzymatic digestion with DNase I (25U/ml) and Collagenase D (1mg/ml) at 37 °C for 20 min under gentle rocking. Tissue suspensions were thoroughly homogenized by gentle and repeated mixing using 10ml pipette to facilitate dissociation of cellular aggregates. Resulting cell suspensions were then filtered through a 40uM cell strainer.

*Flow cytometry analysis and cell sorting*

All samples were analyzed and/or sorted by flow cytometry using a FACSAria III (BD biosciences, San Jose, CA, USA). CD45 (Clone 104) (Biolegend, San Diego, CA, USA), CD45.1 (Clone A20) (BD Biosciences), CD45.2 (Clone 104) (Biolegend), Lineage cocktail (include CD3, B220, Gr1, CD11b, Ter119; Biolegend), cKit (Clone 2B8, Biolegend), Sca1 (Clone D7, Biolegend), EPCR (CD201) (Clone eBio1560, eBioscience, San Diego, CA, USA), CD150 (Clone mShad150, eBioscience), CD48 (Clone HM48-1, Biolegend), and CD31 (Clone 390, BD biosciences) antibodies were used.

*Complete blood counts and in vitro assays*

Peripheral blood was obtained from the facial vein via submandibular bleeding, collected in an EDTA tube, and analyzed using a Hemavet 950FS (Drew Scientific).

For transwell co-culture experiments, $2 \times 10^4$ Lineage negative ($Lin^-$) marrow cells were seeded at the bottom and $1 \times 10^4$ $CD41^+$ MKs seeded on the top inside the transwell. Cells were cultured in StemSpan serum-free expansion medium (SFEM) containing recombinant mouse SCF (100ng/mL) and recombinant human TPO (100ng/mL) (all from Stem Cell Technologies, Vancouver, BC, Canada), as well as 1% antibiotic-antimycotic. Fresh SFEM with cytokines was supplied on day 4 of culture. On day 8, cells were counted and mouse methylcellulose complete media (Methocult GF M3434, Stem Cell Technologies) were used to assay hematopoietic colony formation of post-culture $Lin^-$ cells.

Primary murine lung endothelial cells (ECs) were isolated and cultured as previously described[1]. EC tube formation assay was performed as a measure of angiogenesis *in vitro*. Matrigel® matrix (10mg/ml, Corning Inc., Corning, NY) were thaw overnight at 4°C and kept on ice until use. 150ul Matrigel per well was added to pre-chilled 48-well culture plate. After gelation at 37°C for 30minutes, gels were overlaid with $6 \times 10^4$ primary murine lung ECs (passage 3-4) in 300ul of complete EC medium with SFEM, or Pf4/FF1 MKCM, or control MKCM (all at 1:1 volume ratio). Tube formation was inspected after a period of 2, 4, 6, and 8hrs and images were captured with a phase-contrast microscope (AMEX-1200, AMG, Bothell, WA). The quantification of the capillary tube formation was performed using the ImageJ® software (National Institute of Health, Bethesda, MD) by counting the number of tubes and nodes (branch points) in 4 non-overlapping areas at x40 magnification in two duplicate wells.

*Tissue clearing using modified Murray's clear.*

Immunostained tissues were washed in PBS and dehydrated in a methanol dehydration series (60% methanol for 10 min, then 100% methanol for three 10 min rotations) followed by a 2hr incubation in 100% methanol with several changes of fresh methanol. The methanol was then exchanged with BABB (1:2 Benzyl Alcohol: Benzyl Benzoate). The BABB was cleaned of peroxides by adding 10 g of activated aluminium oxide (Sigma) to 40 ml of BABB and rotating for at least 1 h, then centrifuging at 2000g for 10 min to remove the suspended aluminum oxide particles. The tissues were incubated in BABB for 1-2 days with several exchanges of fresh BABB. Half tibia samples mounted in BABB between two coverslips and sealed with silicone (premium waterproof silicone II clear, General Electric).

*Transcriptome analysis using RNA sequencing*

Wild-type and JAK2V617F mutant marrow $CD41^+$ MKs were isolated using PE-conjugated anti-mouse CD41 antibody (Biolegend) and EasySep Mouse PE Positive Selection Kit II (Stem Cell Technologies) (with 95% purity as measured by flow cytometry analysis). Total RNA was extracted using the RNeasy mini kit (Qiagen, Hilden, Germany). RNA integrity and quantitation were assessed using the RNA Nano 6000 Assay Kit of the Bioanalyzer 2100 system (Agilent Technologies, CA, USA). For each sample, 400ng of RNA was used to generate sequencing libraries using NEBNext® Ultra™ RNA Library Prep Kit for Illumina® (New England BioLabs, MA, USA) following manufacturer's recommendations. The clustering of the index-coded samples was performed on a cBot Cluster Generation System using PE Cluster Kit cBot-HS (Illumina) according to the manufacturer's instructions. After cluster generation, the library preparations were sequenced on an Illumina platform. Index of the reference genome was built using hisat2 2.1.0 and paired-end clean reads were aligned to the reference genome using HISAT2. HTSeq v0.6.1 was used to count the reads numbers mapped to each gene. Differential gene expression analysis of MKs from young Pf4-cre control mice (n=3), young $Pf4^{+}FF1^{+}$ mice (n=3), old Pf4-cre control mice (n=3), and old $Pf4^{+}FF1^{+}$ mice (n=4), was performed using the DESeq R package (1.18.0). Genes with an adjusted $P$-value < 0.05 found by DESeq were assigned as differentially expressed.

*Cytokine Antibody Array*

A membrane array with capture antibodies was used for simultaneous detection of 62 different mouse cytokines according to the manufacturer's protocol (ab133995, Abcam, Cambridge, MA). Briefly, murine $CD41^{+}$ MKs were isolated using the column-free magnetic EasySep™ system (StemCell Technologies) and lysed in cell lysis buffer. Pooled MK cell lysate from young Pf4-cre control mice (n=4), young $Pf4^{+}FF1^{+}$ mice (n=4), old Pf4-cre control mice (n=4), and old $Pf4^{+}FF1^{+}$ mice (n=4) containing 250ng protein was incubated with the membrane overnight after appropriate blocking of the membrane. After washing, the detection antibody was applied to the membrane and immunoblot images were captured using the FluorChem M imaging system (Protein Simple, San Jose, CA). The intensity of each spot was quantified using the ImageJ® software (National Institute of Health, Bethesda, MD).

1. Zhan H, Ma Y, Lin CH, Kaushansky K. JAK2(V617F)-mutant megakaryocytes contribute to hematopoietic stem/progenitor cell expansion in a model of murine myeloproliferation. Leukemia 2016;30:2332-41.
